# Hebbian learning with elasticity explains how the spontaneous motor tempo affects music performance synchronization

**DOI:** 10.1101/2020.10.15.341610

**Authors:** Iran R. Roman, Adrian S. Roman, Ji Chul Kim, Edward W. Large

## Abstract

A musician’s spontaneous rate of movement, called spontaneous motor tempo (SMT), can be measured while spontaneously playing a simple melody. Data shows that the SMT influences the musician’s tempo and synchronization. In this study we present a model that captures these phenomena. We review the results from three previously-published studies: (1) solo musical performance with a pacing metronome tempo that is different from the SMT, (2) solo musical performance without a metronome at a tempo that is faster or slower than the SMT, and (3) duet musical performance between musicians with matching or mismatching SMTs. These studies showed, respectively, that (1) the asynchrony between the pacing metronome and the musician’s tempo grew as a function of the difference between the metronome tempo and the musician’s SMT, (2) musicians drifted away from the initial tempo toward the SMT, and (3) the absolute asynchronies were smaller if musicians had matching SMTs. We hypothesize that the SMT constantly acts as a pulling force affecting musical actions at a tempo different from a musician’s SMT. To test our hypothesis, we developed a model consisting of a non-linear oscillator with Hebbian tempo learning and a pulling force to the model’s spontaneous frequency. While the model’s spontaneous frequency emulates the SMT, elastic Hebbian learning allows for frequency learning to match a stimulus’ frequency. To test our hypothesis, we first fit model parameters to match the data published in (1) and asked whether this same model would explain the data in (2) and (3) without further tuning. Results showed that the model’s dynamics allowed it to explain all three experiments with the same set of parameters. Our theory offers a dynamical-systems explanation of how an individual’s SMT affects synchronization in realistic music performance settings, and the model also enables predictions about performance settings not yet tested.

**Author summary:** Individuals can keep a musical tempo on their own or timed by another individual or a metronome. Experiments show that individuals show a specific spontaneous rate of periodic action, for example walking, blinking, or singing. Moreover, in a simple metronome synchronization task, an individual’s spontaneous rate determines that the individual will tend to anticipate a metronome that is slower, and lag a metronome that is faster. Researchers have hypothesized the mechanisms explaining how spontaneous rates affect synchronization, but no hypothesis can account for all observations yet. Our hypothesis is that individuals rely on adaptive frequency learning during synchronization tasks to adapt the rate of their movements and match another individual’s actions or metronome tempo. Adaptive frequency learning also explains why an individual’s spontaneous rate persists after carrying out a musical synchronization task. We define a new model with adaptive frequency learning and use it to simulate existing empirical data. Not only can our model explain the empirical data, but it can also make testable predictions. Our results support the theory that the brain’s endogenous rhythms give rise to spontaneous rates of movement, and that learning dynamics interact with such brain rhythms to allow for flexible synchronization.

## Introduction

Humans can effortlessly sing or walk showing a spontaneous singing tempo or walking pace. In everyday life, however, they also synchronize with external signals, like singing with a pre-recorded song or marching in a parade with other individuals. Humans can adapt the frequency of their actions to align with a common tempo kept by others. This requires perception-action coordination (PAC) [41], involving communication between the brain’s sensory and motor areas [47]. Nonetheless, how an individual’s spontaneous rate of action affects PAC is still an open question [64].

Before reviewing previous research, we wish to clarify some terminology. In the existing literature, stimulus timing (i.e. a metronome) is described in terms of inter-onset-intervals (IOIs) measured in (milli)seconds. Similarly, “musical tempo” is sometimes described as the time between beats, or the inter-beat interval (IBI). In contrast, in the music literature the “tempo” is a rate (bpm: beats per minute), or frequency, not a time period. The relationship between a frequency *f* (Hz or cycles per second) and a time period *T* (seconds) is 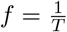. Because we will present a model with an explicit frequency term, when we talk about the SMT (spontaneous motor tempo) and the SPR (spontaneous performance rate) we will use units of Hz. On the other hand, when we talk about the SMP (spontaneous motor period) and the spontaneous performance period (SPP) we will use units of milliseconds. We hope that by acknowledging these historical misnomers we can avoid confusion when trying to understand our model. Table 1 shows a summary of the terms with corresponding abbreviations and units.

**Table 1.**
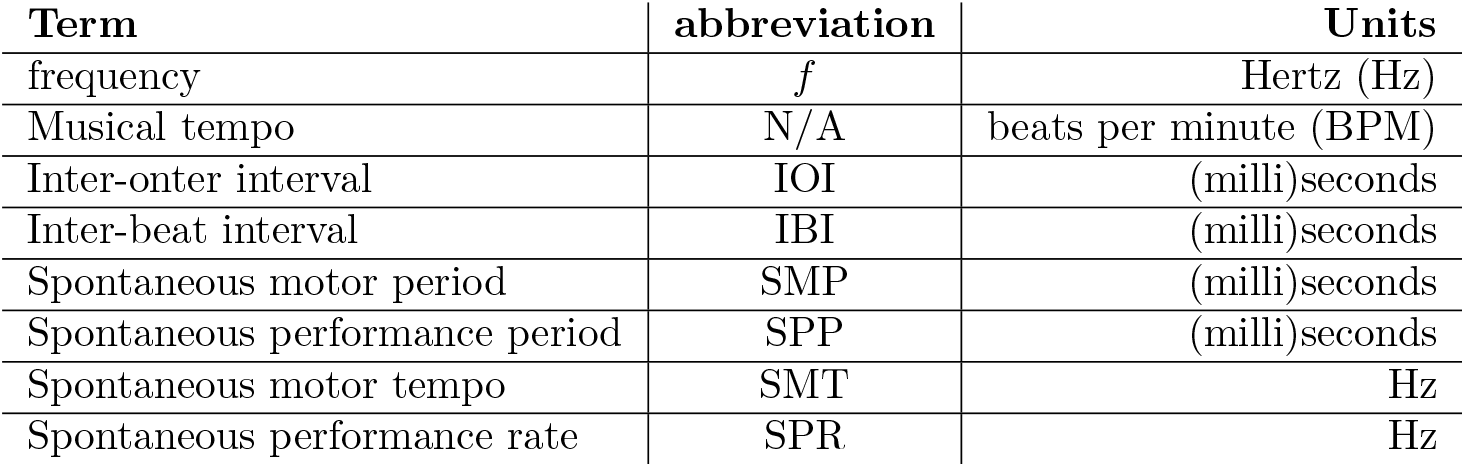
Terminology for periodic phenomena

| Term | abbreviation | Units |
| --- | --- | --- |
| frequency | $f$ | Hertz (Hz) |
| Musical tempo | N/A | beats per minute (BPM) |
| Inter-onset interval | IOI | (milli)seconds |
| Inter-beat interval | IBI | (milli)seconds |
| Spontaneous motor period | SMP | (milli)seconds |
| Spontaneous performance period | SPP | (milli)seconds |
| Spontaneous motor tempo | SMT | Hz |
| Spontaneous performance rate | SPR | Hz |

The SMT can be measured by asking an individual to spontaneously tap or play a simple melody. It tends to be faster in early childhood compared to adulthood [37] and slower in adult musicians (*∼*2.5 Hz on average) compared to non-musicians (*∼*3.3 Hz on average) [9, 52]. The mean asynchrony (MA) can be measured in a synchronization task with a metronome where the average time difference between an individual’s actions and the metronome is calculated [44, 46]. Recent studies have investigated how the SMT and the MA are related. In one study, musicians performed a melody while synchronizing with a metronome faster or slower than their SMT. Results showed that the MA grew as a function of the difference between the metronome tempo and the musician’s SMT. The MA tended to be positive (musicians lagged) when metronome was faster than the SMT, and negative (musicians anticipated) when the metronome was slower than the SMT. Another study looked at musicians performing a melody without a metronome, starting at a tempo different to the SMT. Results showed a tendency to slowly drift back to the SMT [64], an observation also reported in other studies [37, 62]. A different study looked at duet musical performances where pairs of musicians had matching or mismatching SMTs. Matching duets showed smaller MAs compared to mismatching duets. Each musician’s SMT was remeasured after the performance, revealing that the duet synchronization task did not alter each musician’s SMT [63].

These studies highlight the relationship between the SMT, and the MA. However, currently there only exist models that independently explain the underlying mechanisms of SMT and the MA. The SMT has been hypothesized to originate from central pattern generators in the nervous system [16, 30, 35, 52] and motor resonance governed by anatomical properties like body and limb length [13]. The MA has been explained via delayed recurrent feedback in central-peripheral communication between the auditory and motor systems [2, 50, 55] and under- or over-estimation of IOI lengths [33]. A potential way to jointly explain the SMT and the MA could be with non-linear oscillators. While the SMT is equivalent to an oscillator’s natural frequency [26, 27, 37], the MA is analogous to the phase difference between an oscillator and a sinusoidal stimulus once they have reached steady-state phase-locked synchronization. This difference shrinks as the oscillator’s natural frequency and the stimulus frequency become closer [21, 22]. Moreover, if the stimulus ceases the oscillator will return to its natural frequency [21, 22].

This study presents a model able to explain behavioral data that relates the SMT and MA. We use the normal form of an Andronov-Hopf bifurcation [18] as described by Large et al. [28]:

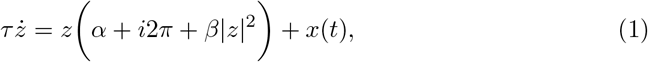

where *z* is the complex-valued state of a non-linear oscillator, *α* and *β* control its dynamics, 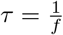 determines its rate (*f* is frequency in Hz), and *x*(*t*) is an external stimulus ^1^. Our model hypothesizes that the SMT originates from central pattern generators [30, 53] and is simulated by Eq (1), which we use in one of its parameter regimes (the supercritical branch of Hopf) due to its limit-cycle properties [28]. However, since Eq (1) can only synchronize with stimulus frequencies close to *f*, its ability to explain both the SMT and the MA is limited [21, 22]. This is clearly distinct from the synchronization abilities of humans, who can synchronize with stimuli tempi that are relatively far away from their SMT [53]. To address this issue we equip Eq (1) with a frequency learning rule.

The SMT could be described as an attractor state that pulls human adaptive synchronization to a rate of activity for optimal energy use [37, 53, 56]. Righetti et al. described *dynamic Hebbian learning*, allowing an oscillator to adapt its frequency and match a frequency component present in a stimulus [49]. Consistent with Righetti et al. we call this mechanism “Hebbian learning” due to its similarities with correlation-based adaptation in neural networks [20], but we acknowledge that the timescale of *dynamic Hebbian learning* is much faster than the slow and long-term changes usually associated with “Hebbian learning”. However, unlike humans [53], Righetti’s oscillators do not return to the initial spontaneous frequency after stimulation ceases. This issue has been addressed by other modeling studies that added elasticity to frequency and phase adaptation models [10, 23, 51]. For the specific case of the Andronov-Hopf bifurcation, Lambert et al. added linear elasticity [23] to Righetti’s equation, resulting in adaptive frequency learning with a constant pull to a rate of activity *ω*_0_:

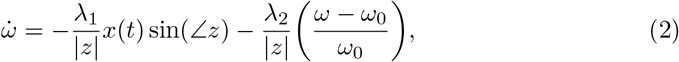

where 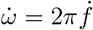 (in radians per second), *ω*_0_ is a fixed “spontaneous frequency” that can simulate the SMT, *λ*_1_ is the frequency learning rate, and *λ*_2_ is the elasticity strength pulling *ω* to *ω*_0_. Combined, Eq (1) and Eq (2) result in an oscillator that can learn the stimulus frequency, but with a force constantly pulling to an otherwise “spontaneous frequency” of activity.

We propose ASHLE (Adaptive Synchronization with Hebbian Learning and Elasticity), a model that builds upon Eq (1) and Eq (2) to explain the underlying neural mechanisms relating the SMT and the MA (ASHLE is mathematically described in the methods section). ASHLE uses two oscillators, “sensory” and “motor”, each simulating the excitatory-inhibitory neural dynamics in sensorimotor networks [16, 28]. While the “sensory” oscillator simulates entrainment of auditory-premotor networks [7, 14, 15, 42], the “motor” one simulates entrainment of motor networks and actions in PAC [12, 29, 60]. In ASHLE, a stimulus can drive and entrain the “sensory” oscillator, which in turn entrains the “motor” one. ASHLE models the SMT as originating from central pattern generators that act as an attractor state for neural and behavioral activity [35]. Therefore, the “motor” oscillator is constantly pulled to a fixed frequency term that simulates the SMT, resulting in an MA between ASHLE’s “motor” oscillator and the stimulus. Behavioral data shows that humans can synchronize with an ongoing stimulus [53], but that they have a tendency to slowly return to their SMT [64]. ASHLE simulates these two timescales by strongly pulling the “motor” oscillator to the SMT, and weakly pulling the “sensory” oscillator to the instantaneous frequency of the “motor” oscillator. These mechanism are the reason why ASHLE uses two oscillators in the first place, because without it the timescales observed in human data would not be possible (see results for experiment 2 and its parameter analysis in the methods section).

Table 2 presents an overview of the most important ASHLE parameters. ASHLE’s oscillators share *α* and *β* to show spontaneous limit-cycle activity. They also share the frequency learning rate *λ*_1_, which controls the *dynamic Hebbian learning* allowing the “sensory” and “motor” oscillators to entrain with a stimulus. *f*_*m*_, the dynamic frequency of the “motor” oscillator, is pulled to a spontaneous rate of activity *f*_0_ with a *λ*_2_ strength. In contrast, *f*_*s*_, the dynamic frequency of the “sensory” oscillator, is weakly pulled, with strength *γ*, to *f*_*m*_.

**Table 2.** ASHLE parameters and function

| Parameter | Function | Value |
| --- | --- | --- |
| $\alpha$ | bifuraction parameter | 1 |
| $\beta$ | non-linear damping | -1 |
| $\lambda_1$ | *dynamic Hebbian learning* | 4 |
| $\lambda_2$ | Elastic pull to $f_0$ | 2 |
| $\gamma$ | Weak elastic pull to $f_m$ | 0.02 |
| $f_0$ | ASHLE’s spontaneous frequency | matches human data |

In this study we use ASHLE to simulate the dynamics of the MA and the SMT.

Fig 1 describes the three previously-published behavioral tasks that we simulate [53, 63, 64]. If ASHLE can explain these dynamics, it would be the first dynamical systems model that systematically captures the relationship between the SMT and the MA as observed in behavioral data. Our results also include simulations that yield predictions that can be tested empirically in future behavioral studies. It is worth noting that ASHLE only simulates the musical beat in a musical performance and not any other spectral features like pitch, harmony or melody content.

**Fig 1.**
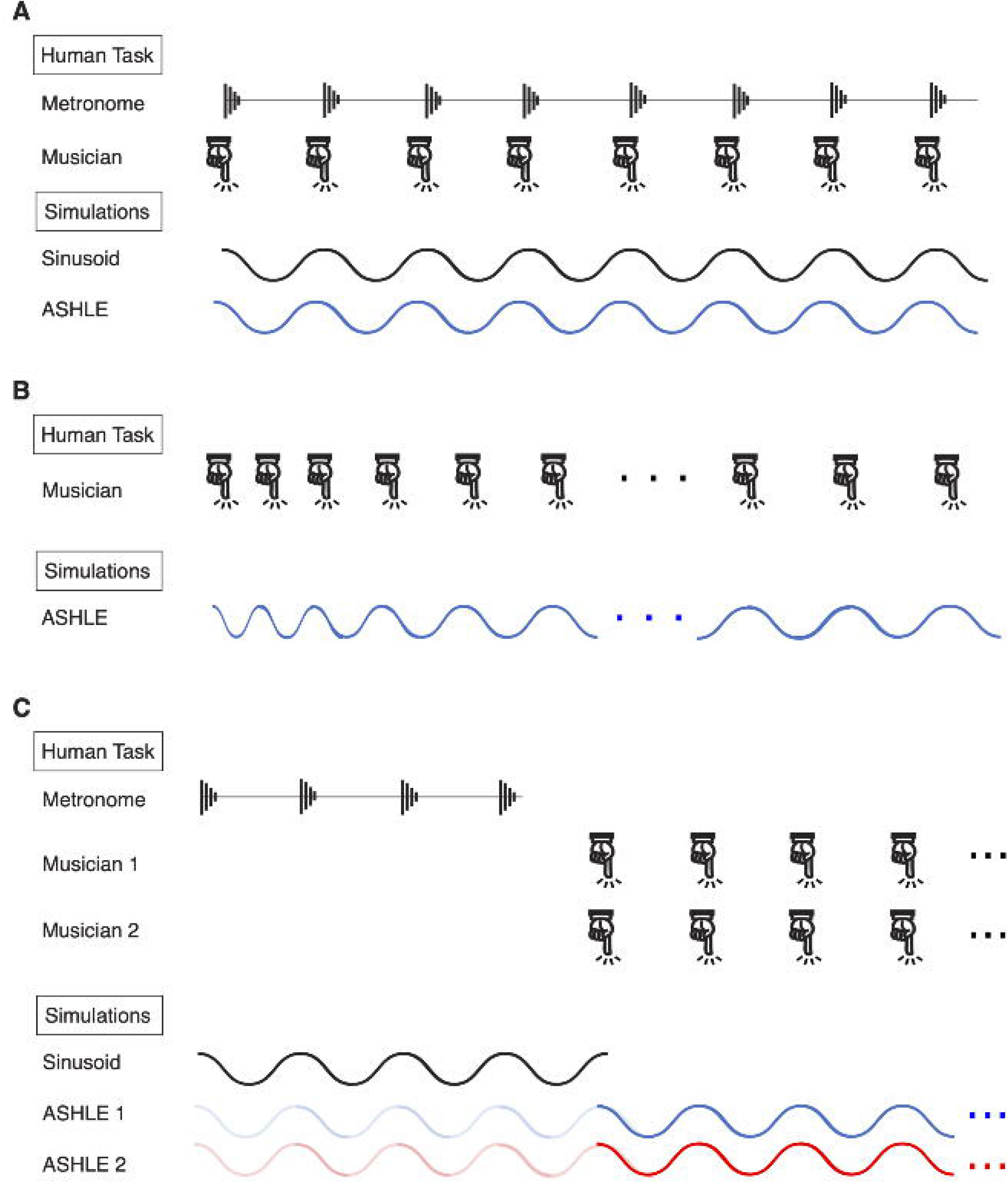
Illustration of the musical tasks and corresponding simulation experiments. (A) The task simulated in experiment 1, in which a musician plays a simple melody with a metronome (top). Illustration of our simulation, in which ASHLE synchronizes with a sinusoidal stimulus (bottom). (B) The task simulated in experiment 2, in which a musician plays a simple melody, without a metronome (top). This specific example shows a performance tempo that periodically became slower due to the musician’s tendency to return to the SMT. Illustration of our simulation, in which ASHLE oscillates without a sinusoidal stimulus and returns to its *f*_0_ (bottom). (C) The task stimulated in experiment 3, in which pairs of musicians played a simple melody together after hearing four pacing metronome clicks (top). Illustration of our simulation, in which two ASHLE models synchronize with four cycles of a pacing sinusoidal stimulus (greyed-out blue and red lines), and then stimulate each other without the sinusoidal stimulus (solid blue and red lines) (bottom).

## Experiment 1: Solo music performance with a metronome tempo different than the SMT

### Results

We simulated the solo task by Scheurich et al. [53] consisting of performance of a simple melody paced by a metronome (Fig 1A). Their experiment had four different experimental conditions: metronome period 30% shorter, 15% shorter, 15% longer, and 30% longer than the musician’s SMP. Fig 2A shows the asynchrony (specifically the “mean adjusted asynchrony”, see methods section for details) between musician and metronome, which was positive (negative) when the metronome period was shorter (longer) than the musician’s SMP. The asynchrony grows as a function of the difference between SMP and metronome period.

**Fig 2.**
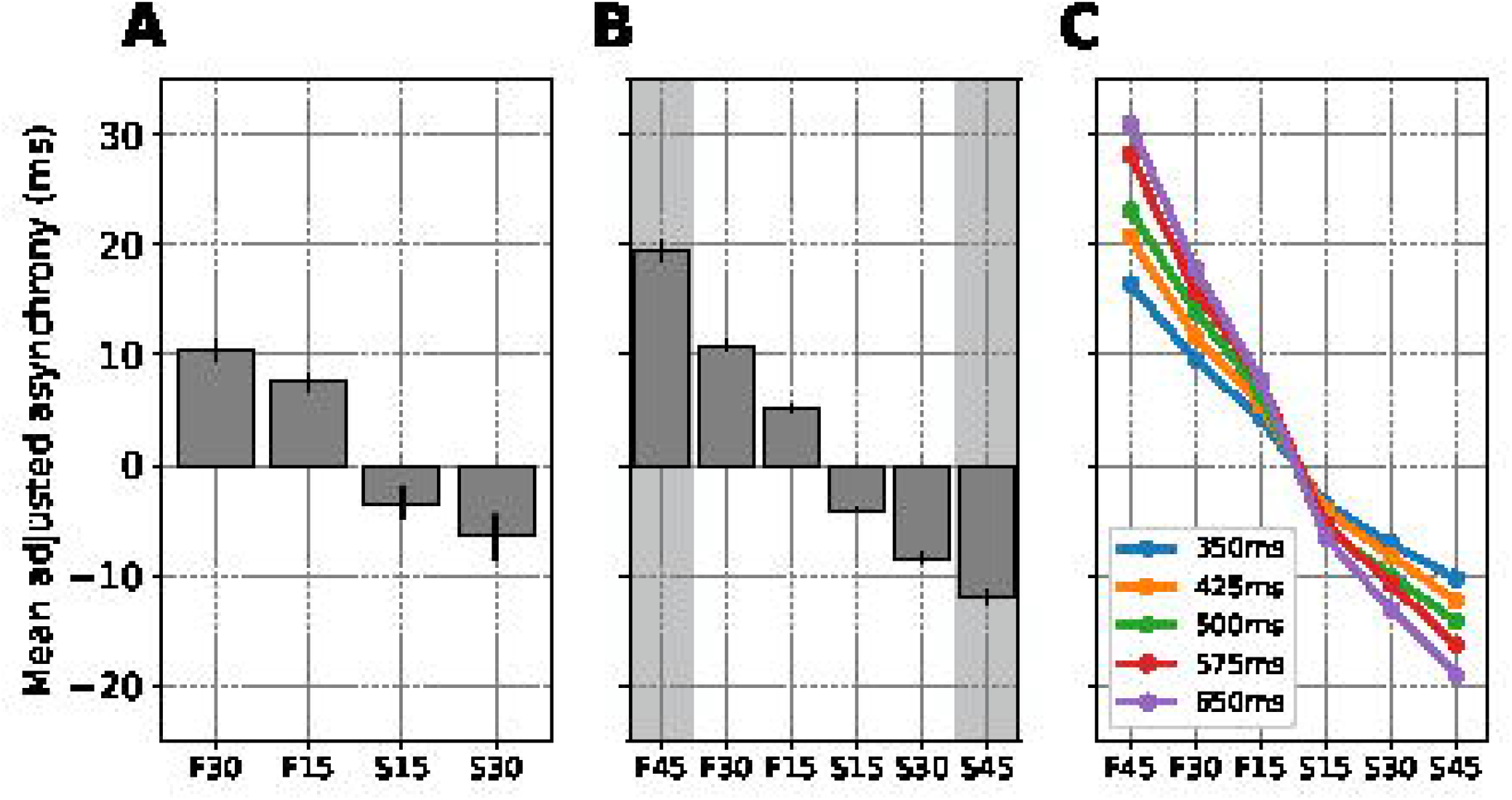
Simulation of the MA between a musician’s beat and a metronome beat with a period shorter or longer than the musician’s SMP during solo musical performance. (A) The mean adjusted asynchrony (and standard error; N=20) between the musician beat and metronome beat during performance of a simple melody in four conditions: metronome period 30% shorter (F30), 15% shorter (F15), 15% longer (S15), and 30% longer (S30) compared to the musician SMP. The x-axis shows “F” and “S” labels as originally used by Scheurich et al. [53] to describe a “faster” and “slower” metronome compared to the SMT. (B) Our simulation results showing the mean adjusted asynchrony (and standard error; N=20) between ASHLE and a sinusoidal stimulus in six conditions: stimulus period 45% shorter (F45), 30% shorter (F30), 15% shorter (F15), 15% longer (S15), 30% longer (S30), and 45% longer (S45) than the period of ASHLE’s *f*_0_. The shaded bars represent predicted measurements for data that has not been collected yet from musicians. (C) Mean adjusted asynchrony predictions when ASHLE models with different *f*_0_ periods synchronize with a stimulus period that is 45% shorter (F45), 30% shorter (F30), 15% shorter (F15), 15% longer (S15), 30% longer (S30), or 45% longer (S45).

A complex-valued sinusoidal stimulus *x* = exp (*i* 2 π *f*_*x*_ *t*) simulated the metronome (*f*_*x*_ is the stimulus frequency in Hz). We hypothesized that ASHLE’s frequency learning, controlled by *λ*_1_, will allow its “sensory” and “motor” oscillators to synchronize with an arbitrary stimulus period. We also expected to see an asynchrony between ASHLE’s “motor” oscillator and the stimulus due to *λ*_2_ pulling the “motor” oscillator to *f*_0_. We simulated 20 different ASHLE models, each with a unique *f*_0_ value that matched a musician’s SMT, as measured by Scheurich et al. [53]. Fig 2B shows that ASHLE can explain the behavioral data, and shaded bars show ASHLE’s prediction for the same group of musicians performing with metronome periods 45% shorter or longer than their individual SMP. Fig 2C also makes behavioral predictions, by breaking down how different ASHLE models with different *f*_0_ synchronize with a metronome period that is 45%, 30%, and 15% shorter or longer than the period of *f*_0_.

### Discussion

Fig 2B showed that our model captures the synchronization dynamics observed in Fig 2A. Specifically, when synchronizing with stimuli slower than *f*_0_, ASHLE showed a negative MA, and a positive MA when synchronizing with stimuli faster than *f*_0_. This is explained by the asymmetry of entrained *f*_*s*_ and *f*_*m*_ around *f*_0_, and the elastic pull of *f*_*m*_ to *f*_0_ by *λ*_2_. In other words, while ASHLE can entrain with the stimulus, the MA results from the pull that makes *f*_*m*_ always be “shy” from perfectly matching *f*_*x*_, the stimulus frequency. It is interesting to note that a closer look at Fig 2A reveals mean adjusted asynchrony magnitudes that are slightly asymmetric between faster and slower stimuli, which implies frequency scaling of PAC. ASHLE has explicit mechanisms that allow it to explain this asymmetry. First, ASHLE has frequency-scaling (see Eqs (3b), (3d)) previously used on Andronov-Hopf oscillator models [21, 28]. Second, ASHLE’s elastic pull is an exponential function (see Eq (3d)). This means that when synchronizing with faster stimuli the pull to its spontaneous frequency is exponentially larger compared to synchronization with slower stimuli. Together, these two mechanisms cause higher *f*_*s*_ and *f*_*m*_ values to amplify the learning rate (*λ*_1_) and the elastic force (*λ*_2_), and lower *f*_*s*_ and *f*_*m*_ values to shrink them.

ASHLE also made testable predictions shown in Fig 2B and Fig 2C. While Fig 2B contains predictions if the same group of musicians were to carry out the task synchronizing with a metronome period 45% shorter or longer than the SMP, Fig 2C contains predictions at the individual musician level, simulating the mean adjusted asynchronies that musicians with different SMP would produce when performing a melody with various metronome tempi faster or slower than their SMT. These predictions can be empirically tested to further validate ASHLE’s behavior or better tune its parameters.

## Experiment 2: Unpaced solo performance with a starting tempo different than the SMT

### Results

Experiment 1 showed that ASHLE can explain the MA as captured by Scheurich et al. [53]. In this second experiment, we used ASHLE to simulate the data by Zamm et al. [64], who studied what happens when musicians perform a simple melody unpaced, but starting at a tempo different than the SMT (Fig 1B). They tested four experimental conditions: starting performance tempo fast, faster, slow, and slower than the SMT. Fig 3A shows their results, which for each condition is the average of the slope (specifically the “mean adjusted slope”, see methods section for details) across musicians. Their data shows that when musicians started at a tempo faster than the SMT the mean adjusted slope was positive (musicians slowed down; consecutive IBIs became longer), and when musicians started at a tempo slower than the SMT the mean adjusted slope was negative (musicians sped up; consecutive IBIs became smaller). In general, musicians showed a tendency to return to their SMT [64].

**Fig 3.**
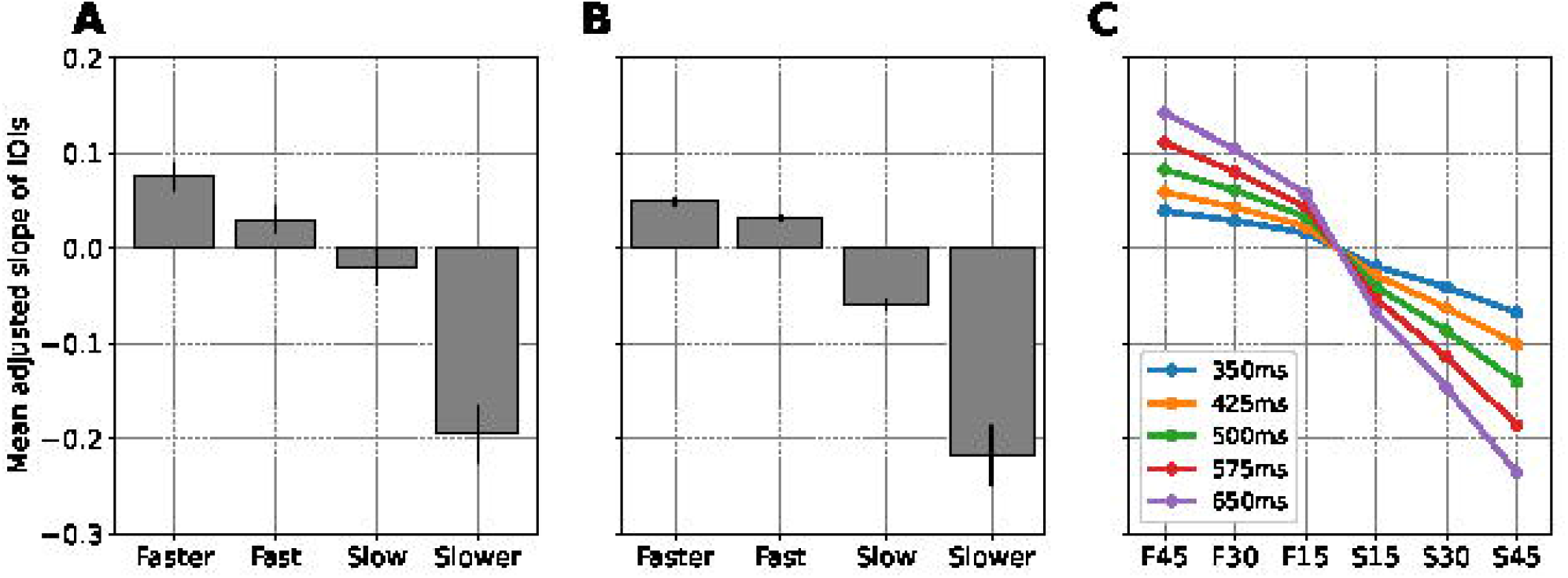
Simulation of the slope between consecutive IBIs when an unpaced musician performs a melody starting at a tempo that is different than the SMT. (A) The mean adjusted slope of consecutive IBIs (and standard error; N=24) when solo musicians perform a simple melody starting a tempo that is fast, faster, slow, or slower compared to their SMT. (B) Our simulations showing the mean adjusted slope of consecutive IBIs (and standard error; N=23) when ASHLE oscillates, without a stimulus, starting at a frequency that is fast, faster, slow, or slower compared to its *f*_0_. (C) Adjusted slope predictions when different ASHLE models with different *f*_0_ oscillate without stimulation, starting with a period that is 45% shorter (F45), 30% shorter (F30), 15% shorter (F15), 15% longer (S15), 30% longer (S30), or 45% longer (S45) compared its to the period of its *f*_0_. For consistency with predictions made in experiment 1, here we also use the F and S on the x-axis

We used the same set of ASHLE parameter values as in experiment 1. The only differences were that stimulus was nullified, ASHLE’s *f*_0_ was dictated by a different set of human SMTs, and initial conditions for *f*_*s*_(0) = *f*_*m*_(0) matched human data by Zamm et al. [64]. We hypothesized that the “motor” oscillator will show a tendency to return to ASHLE’s *f*_0_ as a result of two mechanisms: *λ*_2_ pulling to *f*_*m*_ to *f*_0_, and *γ* pulling *f*_*s*_ to *f*_*m*_. Fig 3B shows that ASHLE can explain the human data observed in Fig 3A. Additionally, this experiment also yielded predictions of human behavior not yet tested shown in Fig 3C..

### Discussion

Our model was also able to simulate the human data by Zamm et al. [64] where musicians performed a melody without a metronome. ASHLE’s approximation of the data is not perfect but, keeping in mind that the parameters *λ*_1_ and *λ*_2_ were optimized with the data in experiment 1, it is close. Hence, results also show the generalization of ASHLE across different datasets and tasks. Since there was no stimulus in this second experiment (*F* = 0), *γ* = 0.02 was optimized to control how quickly ASHLE returns to its *f*_0_.

Experiment 2 highlights potential mechanisms that explain how the SMT influences tempo maintenance by a musician during a solo performance. The small *γ* value is explained by the behavioral data in Fig 3A, showing that musicians tended to very slowly return to their SMT. In other words, *γ* represents a musician’s slow tendency to return to their SMT in the absence of stimulation. We predict that non-musicians will show a faster tendency to return to their SMT since musical training prepares musicians to maintain an arbitrary tempo throughout a musical performance [11, 54]. Future experiments could control ASHLE’s *γ* to explain non-musician data.

ASHLE also made testable predictions shown in Fig 3C at the individual musician level when performing a melody starting at various tempi that are faster or slower than the SMP. These predictions can be empirically tested to further validate ASHLE’s dynamics.

## Experiment 3: Duet musical performance between musicians with matching or mismatching SMTs

### Results

Experiments 1 and 2 showed that the same ASHLE model can simulate two tasks carried out by solo musicians. In this third experiment, we used ASHLE to simulate another task by Zamm et al. [63] showing how musician duets perform a simple melody four consecutive times (Fig 1C). Musician duets were separated into two experimental groups: pairs with matching SMTs and pairs with mismatching SMTs. Fig 4A shows their results, which consisted of the mean absolute asynchrony between the beats of the two performing musicians in each experimental group, separately for each of the four melody repetitions. Their results show that the mean absolute asynchrony was smaller between musician duets with matching SMTs.

**Fig 4.**
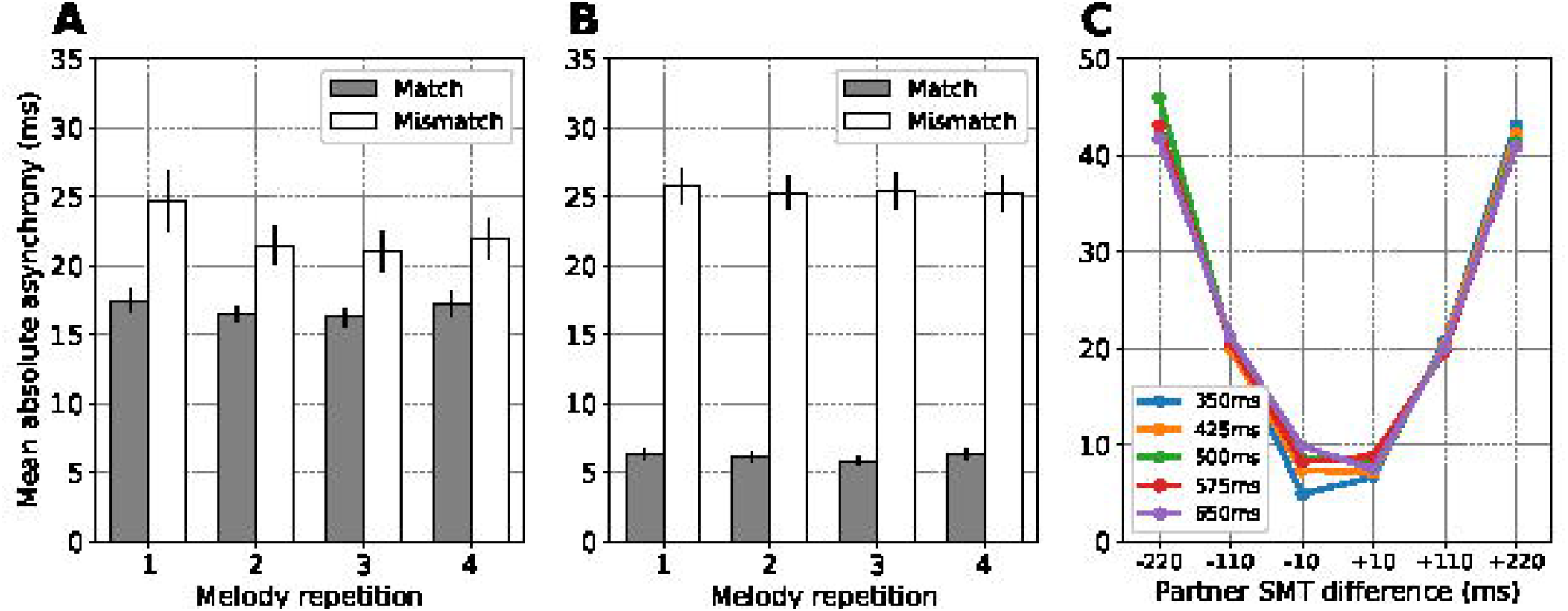
Simulation of the mean absolute asynchrony between two musicians with matching or mismatching SMTs during duet musical performance. (A) The mean absolute asynchrony (and standard error; N=10 per experimental group) between two musicians with matching or mismatching SMTs during performance of a simple melody four consecutive times. (B) Our simulation results showing the mean absolute asynchrony between two synchronizing ASHLE models with *f*_0_ values that are close or far from each other. (C) Mean absolute asynchrony predictions when different ASHLE models (with different *f*_0_ periods) synchronize with another ASHLE model with a *f*_0_ period that is 220ms shorter, 110ms shorter, 10ms shorter, 10ms longer, 110ms longer, and 220ms longer.

In this third experiment we also used the same ASHLE parameters as in experiments 1 and 2. However, pairs of ASHLE models are weakly coupled serving as input to each other. We also we added Gaussian noise to the “motor” oscillator to better match the magnitudes and variances in absolute asynchrony observed in the behavioral data (see model optimization in the methods section). We hypothesized that pairs of ASHLEs will be able to synchronize due to frequency learning. However, the pull of *f*_*m*_ to *f*_0_ will result in asynchrony, which we expect to be smaller between ASHLE pairs with similar *f*_0_. We simulated this task using 20 pairs of ASHLE models with *f*_0_ that matched the 40 musician SMTs measured by Zamm et al. [63]. Fig 4B shows the mean absolute asynchrony observed in our simulations for the same experimental groups and melody repetitions tested by Zamm et al. [63]. Our results are similar to the human data. Additionally, Fig 4C shows musician data predictions by simulating ASHLE duets where one’s *f*_0_ has a period that is 220ms shorter, 110ms shorter, 10ms shorter, 10ms longer, 110ms longer, and 220ms longer than the other. Results in Fig 4C systematically describe how the mean absolute asynchrony between musician pairs grows as a function of the difference between their SMTs.

### Discussion

ASHLE was also able to simulate duet performances (Fig 4A and Fig 4B). To capture the data of experiment 3, it was necessary to add noise to ASHLE’s motor oscillator (see model optimization in the methods section). Without it, ASHLE showed the same qualitative pattern of results, but with a larger difference in magnitude between experimental groups (see Fig 5). When noise was added, ASHLE showed a mean absolute asynchrony around 15ms and 20ms for the duets with matching and mismatching SMTs, respectively. Moreover, the observed standard error in our simulations is similar to the one observed in the human data. This makes sense as the musical performance task has multiple sources of variability, including: two performing musicians receiving feedback from each other, each with variable behavior that will influence the resulting absolute asynchrony between the two.

**Fig 5.**
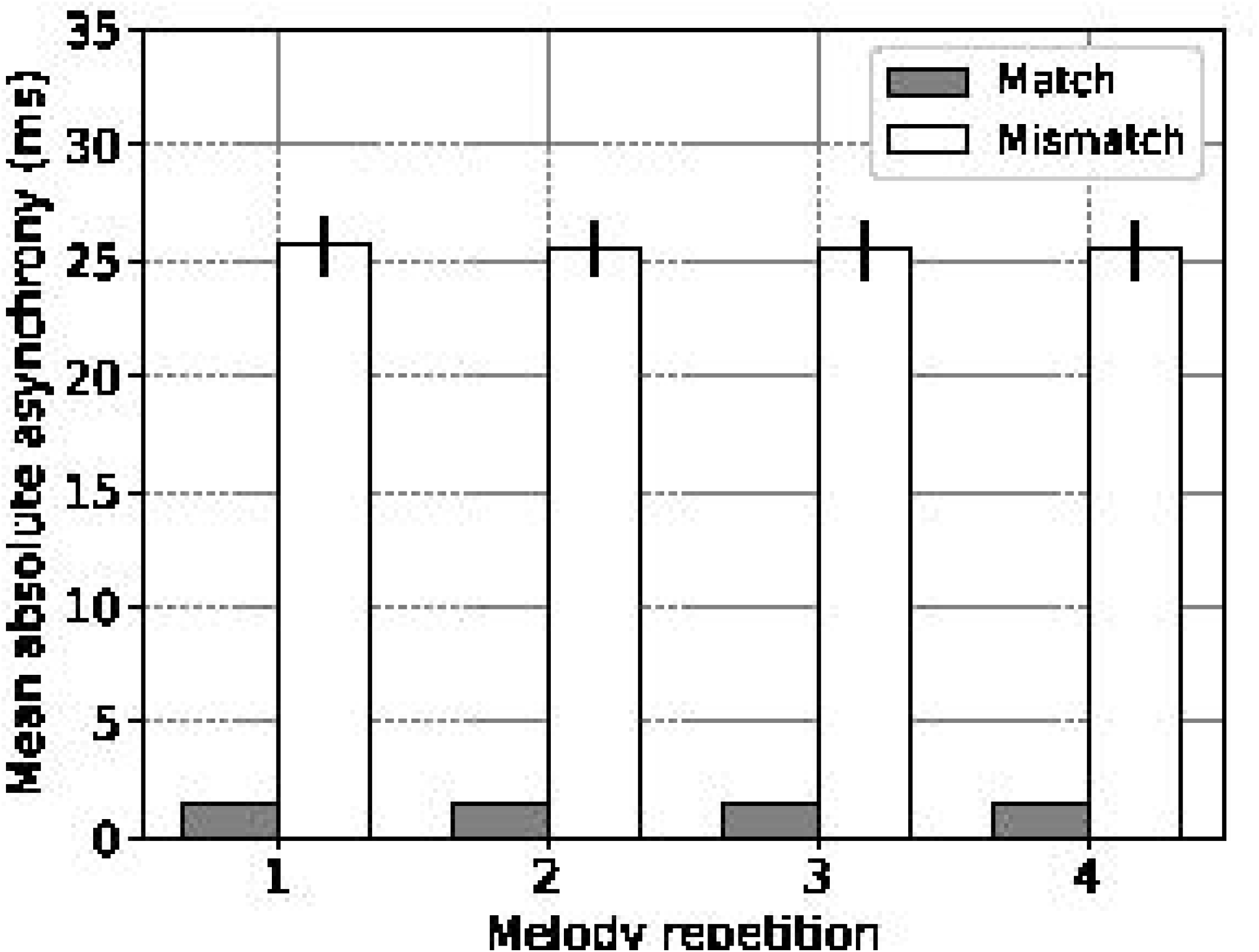
Simulation without noise of the mean absolute asynchrony between two musicians with matching or mismatching SMTs during duet musical performance. Our simulation results showing the mean absolute asynchrony between two synchronizing ASHLE models with matching or mismatching *f*_0_, but no noise added to Eq (3c). The resulting mean absolute asynchronies in this simulation without noise are much smaller compared to the musician data results in Fig 4A. The added noise in Eq (4) improves the model’s results, which are shown in Fig 4B.

To simulate this musical duet task, we also needed smaller connection strength between pairs of ASHLE models (see model optimization in the methods section). This difference is due to the nature of the stimulus. Sinusoidal stimulation of a Hopf oscillator yields stability regimes with phase-locking that widen as a function of stronger forcing [21]. In contrast, two interacting dynamical systems benefit from weaker coupling to exploit emergent resonance higher-order terms (only up to quadratic in ASHLE’s case) [18, 21, 22].

ASHLE also made testable predictions about how musician duets would perform in new experimental manipulations. Fig 4C shows that the mean absolute asynchrony between the two musicians will generally vary as a function of the difference between the musician’s SMTs, and also as a function of the specific SMT value in either of the two musicians. These predictions can also be empirically tested to further validate ASHLE.

### General Discussion

We have presented ASHLE, a model able to simulate human synchronization dynamics. ASHLE can explain how a musican’s SMT results in asynchronies with a pacing metronome during a simple melody performance task (Fig 2). It can also explain the rate of tempo change when a solo musician performs a simple melody without a metronome (Fig 3). Two ASHLE models can also be coupled to explain the asynchrony between two musician performing a melody as a duet (Fig 4). ASHLE’s adaptive frequency learning mechanism allows it to generalize across these different tasks while allowing for testable predictions of human data not yet collected (Fig 2B, Fig 2C, Fig 3C, and Fig 4C).

### ASHLE’s elastic frequency learning captures diverse synchronization dynamics

Experiments 1 and 3 showed that the asynchrony between ASHLE and stimulus is the result of the pull of *f*_*m*_ to *f*_0_. Experiment 2 showed that, in the absence of a stimulus, ASHLE can start oscillating at a rate higher or slower than its *f*_0_, but that the same elastic pull progressively causes it to return to *f*_0_ via the slow tendency of *f*_*s*_ to converge with *f*_*m*_. ASHLE’s behavior is consistent with theoretical accounts highlighting that a dynamical system’s natural frequency (i.e. ASHLE’s *f*_0_) is the optimal state for synchronization, even if the system can synchronize at other frequencies [17, 19, 53, 59].

An interesting question is where in the brain these mechanisms of adaptive frequency learning and elasticity occur. ASHLE is a working hypothesis of neuroscientific mechanisms underlying PAC. There exist theories that explain the human SMT as originating from central pattern generators [16, 30, 35, 53] and ASHLE’s *f*_0_ is a parameter and attractor state consistent with this theory. Additionally, research indicates that cortical and subcortical sensorimotor networks synchronize neural activity that reflects the rhythms of a periodic stimulus and PAC [6, 12, 14, 29]. ASHLE’s oscillators show sustained oscillatory activity to simulate entrainment in sensorimotor brain areas when processing a periodic stimulus of an arbitrary frequency [7, 14, 15].

Moreover, ASHLE’s “sensory” and “motor” oscillators show activity synchronized with a stimulus, thus simulating neural entrainment to the musical beat [29]. We also consider the peaks of the “motor” oscillator as an indicator of entrained peripheral effectors (i.e., a finger playing a piano) during a musical performance task. In summary, stimuli entrain ASHLE’s “sensory” oscillator that simulates sensory and premotor neural dynamics. This in turn entrains the “motor” oscillator that simulates motor network dynamics and the transformation into motor commands.

While no consensus exists yet about the interplay of the SMT in PAC, ASHLE is a working model of these hypotheses with mechanisms that explain how a constant pull to the rate of activity of a central pattern generator affects adaptive PAC. Similar models with period and phase correction mechanisms have also been characterized for their flexible and multi-stable phase-locking dynamics [5, 10, 34, 51], and together with ASHLE support our hypothesis of the underlying elastic frequency learning dynamics of PAC.

### ASHLE’s relation to models of the negative mean asynchrony

Our study is part of a larger effort to characterize human PAC dynamics. The mean asynchrony has been of particular interest for existing models of PAC [5, 43, 50, 55]. It has been shown to be negative for stimulus with IOIs greater than 300ms but smaller than 2000ms [36], growing as a function of IOI duration. This phenomenon is known as the negative mean asynchrony (NMA). For IOIs smaller than 300ms, the MA has been reported to be absent or slightly positive [43, 61], indicating that 300ms is the transition point for MA to go from negative to positive values. Existing modeling work has largely explained the NMA independent of the SMT, with models proposing that the NMA is the result of adaptive synchronization by means of phase and period correction rules at phenomenological [58] and neuromechanistic levels [5]. Other models follow the strong anticipation hypothesis [55], explaining that the NMA results from delayed neural communication in the sensorimotor system [50].

While ASHLE showed both positive and negative asynchronies, its parameters were optimized to simulate the “mean adjusted asynchrony” reported in the behavioral data of Scheurich et al. [53]. Therefore, what seems to be an NMA in Fig 2A (negative-valued bars) is in reality something else and ASHLE cannot be considered to be an NMA model. A future direction could be to add delayed-feedback to ASHLE’s oscillators and frequency adaptation rules to allow it to explain the NMA while also accounting for the SMT. We also invite human researchers to collect and report both the SMT and the NMA of individual participants in future behavioral studies.

### ASHLE’s potential to simulate non-musician data and its current lack of variability

Behavioral data shows that musical expertise affects the MA, with musicians showing overall smaller MAs compared to non-musicians [45, 53]. While we optimized ASHLE to explain musician data, its parameters can be controlled to account for non-musician behavior. First, we determined ASHLE’s spontaneous frequency using individual musicians’ SMTs. To simulate non-musician data one would only need to change these values to match a non-musician population. Larger MAs by non-musicians [45, 53] also imply a stronger pull to their SMT, which would translate into ASHLE’s parameters *λ*_2_ and *γ* being larger than the ones we found.

While ASHLE was able to simulate the musician data in experiments 1 and 2, it does so by reaching a steady-state with little or zero variability (see the variance of Fig 2B and Fig 3B). This was a limitation to perfectly simulate the behavioral results since standard errors were overall smaller in ASHLE’s simulations compared to the behavioral data. In empirical studies of PAC by either musicians or non-musicians there exist two sources of variability: (1) variability within each particiant’s behavior across time and (2) variability between the behavior of different particiant. In its current form ASHLE only allowed us to simulate the first one, using different ASHLE models with different spontaneous frequency values. Future research could add gaussian noise to ASHLE’s activity and characterize its ability to better approximate the variance associated with human data. Our study includes an early attempt of this by adding noise to ASHLE’s “motor” oscillator in experiment 3.

### ASHLE’s relation to gradient frequency neural network models

ASHLE builds upon the work by Large et al. [28] and Lambert et al. [23], who proposed Gradient Frequency Neural Networks (GFNNs). Lambert et al. used a bank of oscillators but, in contrast with Large et al., each oscillator had the ability to adapt its frequency to resonate with a dynamic stimulus (musical rhythms in their case).

Lambert et al. also used an elasticity term to pull each oscillator to a fixed central frequency and ensure that oscillators did not overlap in frequency with each other. This allowed them to synchronize their network similarly to Large et al. but using significantly less oscillators.

ASHLE is different in many ways. First, its elastic frequency learning is used to simulate synchronization tasks where asynchronies were observed and explained as a function of the SMT. Second, ASHLE works at the level of synchronization with a musical beat (most music has a beat around *≈*1.5 Hz), a specificity that allows ASHLE to directly process a stimulus using its “sensory” oscillator. Third, ASHLE’s rates of activity in its “sensory” and “motor” oscillators are centered around the SMT (*≈*2.5Hz) and overlap with the delta band of neural oscillation, which has been hypothesized to be critical for the processing of musical rhythms [8]. Therefore, ASHLE is a minimal model that is specific to simulate behavioral. Future research could look into making ASHLE’s more general. We hypothesize, for example, that it will be possible to use a network of oscillators with elastic frequency learning to extract the beat information directly from a “raw” musical signal. If such oscillators are centered around integer ratios of the SMT, the overall oscillatory activity will reflect synchronization with the stimulus but with a constant pull to the SMT.

### Conclusion and Outlook

We have presented a versatile model of human behavior during a musical beat synchronization task. ASHLE was able to explain empirical data from three different human studies, and related human behavior with endogenous rhythms like the SMT. Future work could test whether ASHLE’s adaptive frequency learning and synchronization allow it to track semi-periodic signals, such as speech envelopes.

Furthermore, ASHLE could be used not just to simulate and make testable predictions of human behavior, but also as a tool with which humans can interact in experimental, musical, and therapy settings. ASHLE advances our understanding about how sensory-motor entrainment to stimulus frequencies interacts with the constraints of endogenous rhythms (i.e. the SMT) hypothesized to originate from central pattern generators.

## Methods

### The ASHLE model

Eqs (3a), (3b), (3c), and (3d) show the ASHLE model:

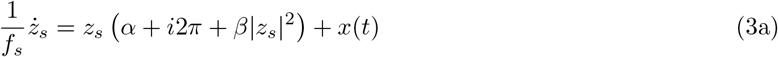

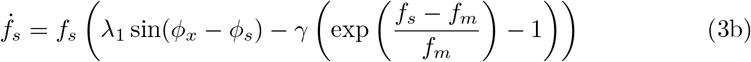

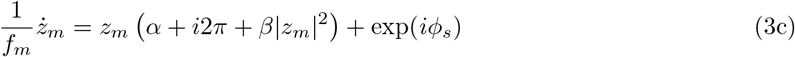

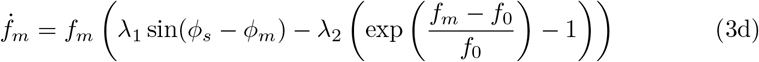

Eqs (3a) and (3c) are oscillators like the one in Eq (1). Eqs (3a) and (3b) have a subscript *s* that stands for “sensory”, while Eqs (3c) and (3d) have a subscript *m* that stands for “motor”. *ϕ* stands for the instantaneous phase of the stimulus (*ϕ*_*s*_), “sensory” (*ϕ*_*m*_) and “motor” (*ϕ*_*x*_) oscillators. In all simulations we run in this study *α* = 1 and *β* = −1 so that the intrinsic dynamics of Eqs (3a) and (3c) are a limit-cycle. In the absence of stimulus (i.e., when *F* = 0), Eqs (3a) and (3c) will show spontaneous and perpetual oscillation [21]. These limit-cycle properties could be achieved with other oscillators. In fact a phase-only (no amplitude term) oscillator could have been used. However, we select Eq (1) due to its neural underpinnings of excitatory-inhibitory oscillation in cortical and subcortical networks [21, 28, 29, 31]. Eqs (3b) and (3d) are the frequency learning equations for Eq (3a) and Eq (3c), respectively. Eq (3b) has a learning term *λ*_1_ sin(*ϕ*_*x*_ *− ϕ*_*s*_) and a slow term 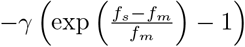. The first one learns the frequency of the external stimulus *x*(*t*), while the second is optimized to weakly pull *f*_*s*_ to *f*_*m*_. This weak pull is important when *F* = 0, allowing Eq (3b) to slowly forget an entrained frequency (see parameter analysis in the next subsection). Eq (3d) entrains to match the frequency of Eq (3b), but *λ*_2_ is optimized to strongly pull *f*_*m*_ to *f*_0_, which is ASHLE’s spontaneous rate of activity.

### Parameter analysis

Previous studies have investigated how Eq 3a synchronizes with a periodic external stimulus in a phase-locked fashion [21]. We focus on analyzing how adaptive and elastic frequency learning, controlled by parameters *λ*_1_ and *λ*_2_, affects synchronization with a periodic stimulus. The dynamics of *γ* will be analyzed in the methods subsection of its relevant experiment (experiment 2).

We set ASHLE’s spontaneous frequency *f*_0_ = 2.5 because the average musician SMP is around 400ms in previous behavioral studies 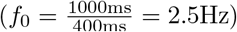 [53, 63, 64]. We analyzed how *λ*_1_ and *λ*_2_ affect the phase-locked asynchrony between the stimulus *x*(*t*) and ASHLE’s *z*_*m*_ when the stimulus period is 45% shorter 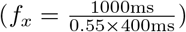 and 45% longer 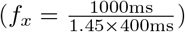 than ASHLE’s *f*_0_ period. We ran fifty-second-long simulations, each with a unique value for *λ*_1_ and *λ*_2_, and *x*(*t*) = exp(*i*2*πf*_*x*_*t*), where *f*_*x*_ is the stimulus frequency in Hz and *t* is time. Initial conditions were *z*_*s*_(0) = *z*_*m*_(0) = 0.001 + *i*0, and *f*_*s*_(0) = *f*_*m*_(0) = *f*_0_. *γ* = 0 because we do not want to study the effect of this small parameter in this first analysis. After each simulation finished, we found the location (in milliseconds) of all peaks in the real part *x*(*t*) and *z*_*m*_ (Fig 6A). Then, we subtracted the location of the peaks of *x*(*t*) from the location of the peaks of *z*_*m*_ and averaged the result to obtain the mean asynchrony in milliseconds. If *x*(*t*) and *z*_*m*_ showed a different number of peaks, that indicated that phase-locked synchronization did not occur between the two. Fig 6B and Fig 6C show the result of this analysis, with black cells incidating that phase-locked synchronization does not occur for certain combinations of *λ*_1_ and *λ*_2_. Not surprisingly, when *λ*_1_ = 0, phase-locked synchronization is never possible. This makes sense since *λ*_1_ is the frequency learning rate that allows ASHLE to adapt its frequency. When *λ*_1_ = 0, phase-locked synchronization can only occur between ASHLE and a stimulus with a frequency that is close to ASHLE’s *f*_0_ [21]. This analysis also revealed that as the value of *λ*_2_ becomes larger, phase-locked synchronization may not be observed because ASHLE’s *f*_*m*_ is strongly pulled to the spontaneous frequency *f*_0_. The size of the asynchrony is modulated by *λ*_1_ and *λ*_2_. As *λ*_1_ becomes larger, the asynchrony tends to decrease and is sometimes close to zero when *λ*_2_ = 0. The opposite occurs when *λ*_2_ grows. This observation reveals that *λ*_1_ and *λ*_2_ work in opposing directions. While *λ*_1_ changes the model’s frequency to match the stimulus frequency, *λ*_2_ pulls ASHLE’s to *f*_0_. Moreover, as ASHLE’s frequency deviates from *f*_0_, *λ*_2_ acts with more strength, while the strength of *λ*_1_ is not directly affected by the difference between *f*_0_ and ASHLE’s frequency. Additionally, when synchronization is observed between ASHLE and the stimulus *x*(*t*), the sign of the asynchrony between *z*_*m*_ and *x*(*t*) is affected by whether *x*(*t*) is faster or slower than ASHLE’s *f*_0_, with a tendency be positive (lagging) and negative (anticipating), respectively.

**Fig 6.**
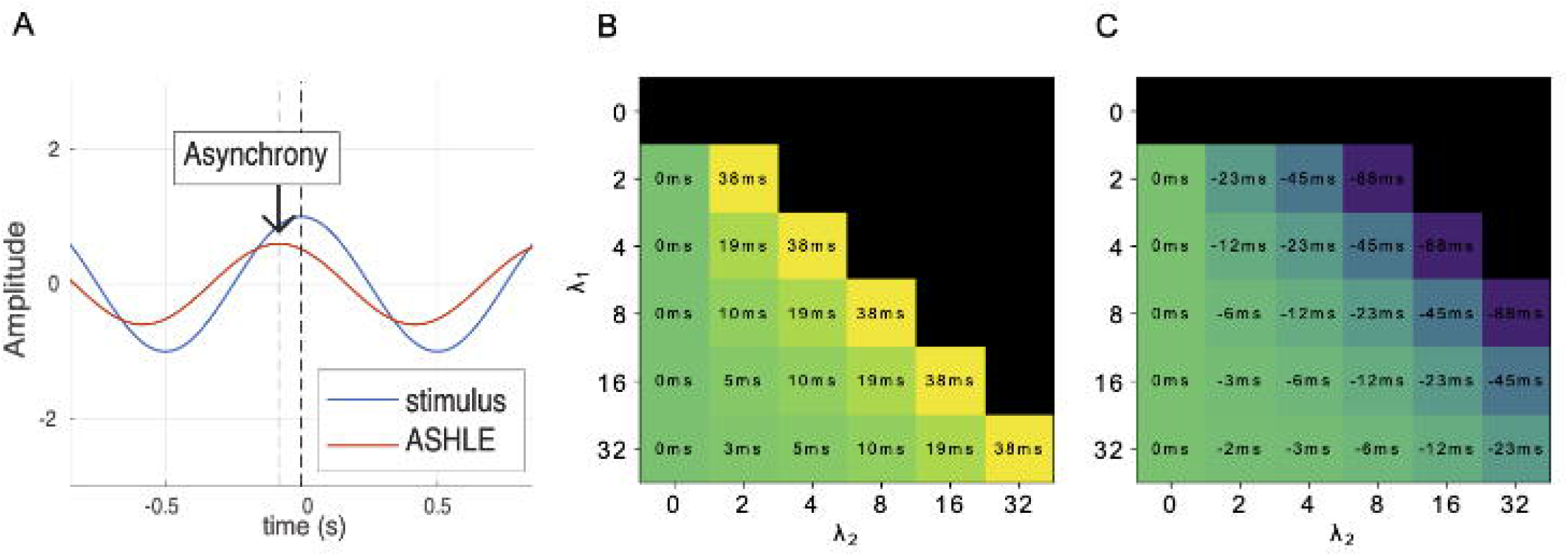
The asynchrony between ASHLE and a sinusoid with period 45% shorter or longer than its spontaneous frequency, as a function of frequency learning and elasticity parameters. (A) Illustration of the asynchrony between ASHLE’s *z*_*m*_ and the sinusoidal stimulus. (B) The asynchrony in milliseconds between ASHLE and a sinusoidal stimulus with a period 45% shorter than ASHLE’s spontaneous frequency, and its change as a function of *λ*_1_ and *λ*_2_. (C) The same analysis but for a sinusoidal stimulus with a period 45% longer than ASHLE’s spontaneous frequency. Black cells indicate *λ*_1_ and *λ*_2_ value pairs for simulations where ASHLE could not synchronize.

## Experiment 1: Solo music performance with a metronome tempo different than the SMT

### Behavioral data for simulation

In the task by Scheurich et al. [53], 20 musicians individually performed a simple melody while synchronizing with a metronome in four experimental conditions: metronome period 30% shorter, 15% shorter, 15% longer, and 30% longer than their SMP. For each metronome rate, participants performed the melody (“Mary had a little lamb”) four consecutive times (32 beats per repetition, 128 beats total). Experimenters measured the mean adjusted asynchrony between the participant beats and the metronome clicks during the middle two melody repetitions (64 beats total). The mean adjusted asynchrony is the MA observed when the musician performs with a metronome tempo different than the musician’s SMT, minus the MA observed when the musician performs with a metronome that matches the SMT. Scheurich et al. [53] used the mean adjusted asynchrony instead of the MA to assume in their analysis that no MA exists between the musician and the metronome that matches the SMT. Fig 2A shows the behavioral data with the mean adjusted asynchrony (average and standard error) observed across all 20 musicians for each experimental metronome tempo condition. Their results showed that the mean adjusted asynchrony had a tendency to be positive when synchronizing with a metronome faster than the SMT (musician actions lagging the metronome), and negative when synchronizing with a metronome slower than the SMT (musicians actions anticipating the metronome). Additionally, the mean adjusted asynchrony grew as a function of the difference between musician SMT and experimental metronome tempo.

### Setup, procedures and measurements

To obtain the musicians SMPs, we overlaid a grid over figure 4 from the paper by Scheurich et al. [53], which showed each musician’s measured SMP (in their original paper, the authors call the SMP as the “SPR”). This allowed us to precisely digitize each musician’s SMP from the original publication. We simulated 20 different ASHLE models, all with the same parameter values (see model optimization below) except for *f*_0_, which had a period that matched the SMP of a different musician. In all simulations, initial conditions were *z*_*s*_(0) = *z*_*m*_(0) = 0.001 + *i*0, 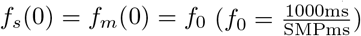. Following the procedure in the musician experiment, each ASHLE synchronized during 128 cycles with a pacing complex-valued sinusoidal stimulus *x*(*t*) = exp(*i*2*πf*_*x*_*t*) where *f*_*x*_ had a period either 30% shorter 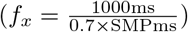, 15% shorter 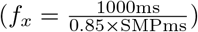, 15% longer 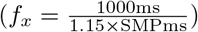, or 30% longer 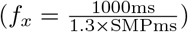 than ASHLE’s *f*_0_ period. We also simulated how ASHLE would synchronize with a stimulus period 45% shorter 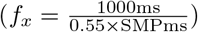 or 45% longer 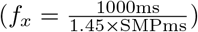 in order to make predictions about how musicians would perform in those additional experimental conditions. After each simulation, we identified the location (in milliseconds) of local maxima in the real part of ASHLE’s *z*_*m*_ and the sinusoidal stimulus. Then, in each simulation we identified the middle 64 peaks for ASHLE and the stimulus and subtracted the location of stimulus peaks from ASHLE peaks to obtain the asynchrony. Averaging these asynchronies in each simulation resulted in the MA for a specific simulation. To obtain the mean adjusted asynchrony, from each MA obtained in the experimental conditions, we subtracted the MA observed when ASHLE synchronized with a sinusoid with a frequency that matched ASHLE’s *f*_0_ (*f*_*s*_ = *f*_0_; after accounting for numerical error we observed that ASHLE had an asynchrony of virtually zero in this condition). We averaged the mean adjusted asynchronies observed across the 20 ASHLE models to obtain the plot in Fig 2B. We also simulated how different ASHLE models with specific *f*_0_ periods (linearly spaced between 350ms and 650ms) would carry out this task when synchronizing with a stimulus period that is 45% shorter 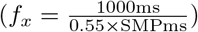, 30% shorter 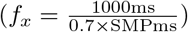, 15% shorter 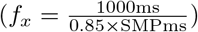, 15% longer 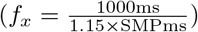, 30% longer 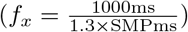, or 45% longer 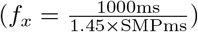 than ASHLE’s *f*_0_ period. Fig 2C shows the results of these simulations, which are predictions of musician data that could be collected to test the accuracy of predictions made by ASHLE.

### Model optimization

We identified the set of ASHLE parameters that result in asynchronies observed in the data by Scheurich et al. [53]. We ran simulations where ASHLE was stimulated by a different stimulus frequency to measure the MA between an ASHLE model with an *f*_0_ = 2.5 (same as our original parameter analysis shown in Fig 6B and Fig 6C) and a sinusoidal stimulus *x*(*t*) = exp(*i*2*πf*_*x*_*t*) with six potential period lengths: 45% shorter 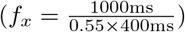, 30% shorter 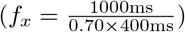, 15% shorter 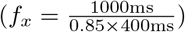, 15% longer 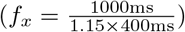, 30% longer 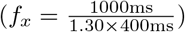, and 45% longer 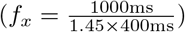. The parameter analysis in the previous section releaved that values for *λ*_1_ between 3 and 5, and *λ*_2_ between 1 and 3 could result in MA values in the range of the mean adjusted asynchrony observed in the study by Scheurich et al. [53] (Fig 6B and Fig 6C). In this analysis we refine our search for *λ*_1_ and *λ*_2_ in this range of values. Subplots in Fig 7 show the asynchrony between *z*_*m*_ and *x*(*t*) for a different stimulus frequency and a pair of parameter values of *λ*_1_ and *λ*_2_. This analysis revealed that the values of *λ*_1_ = 4 and *λ*_2_ = 2 result in the range of MA values observed in humans.

**Fig 7.**
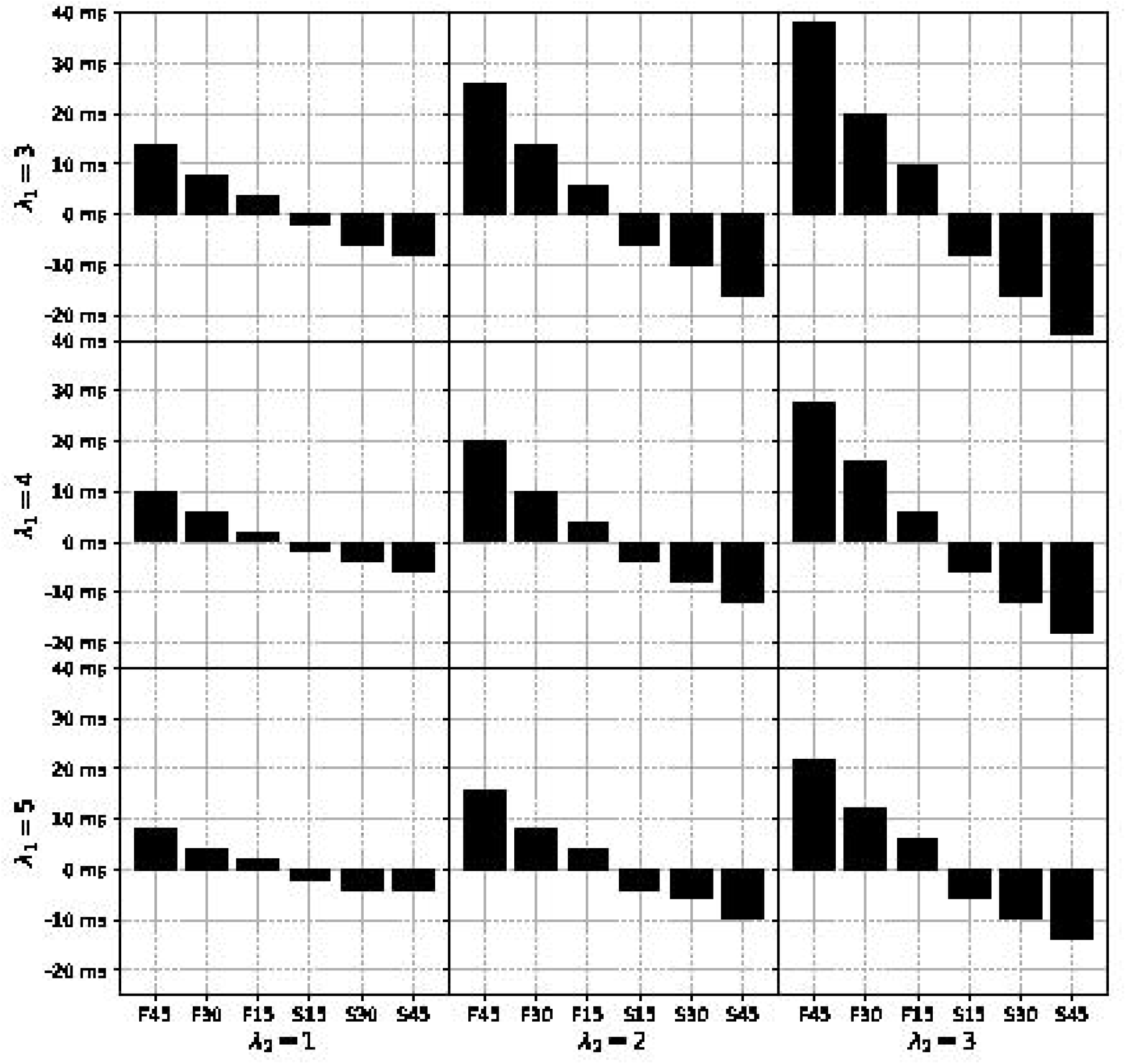
The asynchrony between ASHLE and a sinusoid faster or slower than ASHLE’s *f*_0_ as a function of a narrower range of values for the frequency learning and elasticity parameters. Each cell shows the MA between ASHLE and a sinusoidal stimulus with a period 45% shorter, 30% shorter, 15% shorter, 15% longer, 30% longer, and 45% longer than ASHLE’s 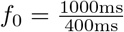 period, for a pair of values for *λ*_1_ and *λ*_2_. The pair of *λ*_1_ = 4 and *λ*_2_ = 2 yield MA values similar to the ones that Scheurich et al. [53] observed in musicians synchronizing with a metronome period 30% shorter, 15% shorter, 15% longer, and 30% longer than the musician’s SMP.

## Experiment 2: Unpaced solo music performance with a starting tempo different than the SMT

### Behavioral data for simulation

In the task by Zamm et al. [64], 24 musicians individually performed a simple melody without listening to a metronome. First they performed the melody at their SMT, and in four other experimental conditions they performed starting at four other spontaneous tempi: fast and slow with respect to SMT, and even faster and slower with respect to the SMT. For each spontaneous initial tempo, musicians performed the melody (“Frere Jaques”) four consecutive times (32 beats per repetition, 128 beats total).

Experimenters measured the IBI across each musician’s entire performance and carried out a linear regression to obtain a slope, which indicated the rate of change across IBIs. Fig 3A shows the behavioral data with the average slope across participants in each initial tempo condition. Results showed that the slope had a tendency to be positive when performances started a spontaneous tempo faster than the SMT (IBIs becoming longer as the performances progressed), and negative when performances started at a spontaneous tempo slower than the SMT (IBIs becoming shorter as the performance progressed).

### Setup, procedures and measurements

To obtain the musicians’ SMP, we overlaid a grid over Figure 1 from the paper by Zamm et al. [64], which showed each musician’s measured SMP (in their original paper, the authors call the SMP as the “SPR”). We also overlaid a grid over Figure 2 (right top panel) from the same paper by Zamm et al. to obtain each musician’s initial rates of performance that were fast, slow, faster, and slower with respect to their SMT. This allowed us to precisely recover each musician’s rates of performances reported in the original study. Using the musicians’ SMT and initial performance tempo values, we simulated 23 different ASHLE models, each with 5 different initial conditions *f*_*s*_(0) = *f*_*m*_(0): (1) matching a musician’s SMT, (2) “fast” or (3) “slow” compared to the SMT, and (4) “faster” and (5) “slower” than the SMT (115 total simulations). We did not simulate the participant with the SMP of 665ms because their “fast” spontaneous tempo was significantly faster than the rest of the participants’ “fast” tempi. ASHLE was not able to show stable activity as a result of this tempo difference.

In all simulations in this experiment there was no stimulus (*F* = 0). Other than *f*_0_ and the initial conditions for *f*_*s*_(0) = *f*_*m*_(0), all simulations shared the same parameter values (see model optimization below). After each simulation, we identified the location (in milliseconds) of local maxima in the real part of ASHLE’s *z*_*m*_. Then, we measured the difference between consecutive peaks to analyze how IBIs change over the course of a simulation. For each simulation we carried out a linear regression over the IBIs to obtain a slope value. Consistent with the methods in the human experiment to obtain the adjusted slope, the slope of each simulation in the control condition where *f*_*s*_(0) = *f*_*m*_(0) = *f*_0_ was subtracted from the slope obtained in the experimental conditions (*f*_*s*_(0) = *f*_*m*_(0)*≠ f*_0_). After accounting for numerical error we observed that ASHLE had slope of virtually zero when *f*_*s*_(0) = *f*_*m*_(0) = *f*_0_. We averaged adjusted slopes across ASHLE models to obtain the plot in Fig 3B. We also simulated how different ASHLE models with specific *f*_0_ values (with period lengths linearly spaced between 350ms and 650ms) would carry out this task when the initial conditions for *f*_*s*_(0) = *f*_*m*_(0) were a period 45% shorter 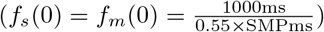, 30% shorter 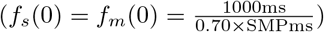), 15% shorter 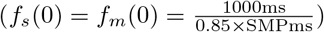, 15% longer 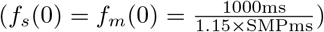, 30% longer 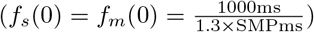, and 45% longer 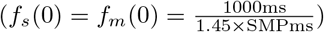 than ASHLE’s *f*_0_ period. Fig 3C shows the results of these simulations, which are predictions of musician data that could be collected to test the accuracy of predictions made by ASHLE.

### Model optimization

We identified the parameter *γ* that results in ASHLE’s return to *f*_0_ when there is no stimulus present (*F* = 0) and the initial values of *f*_*s*_(0) = *f*_*m*_(0) ≠ *f*_0_. To optimize ASHLE we use the same setup as in our original parameter analysis with the exception that initial conditions for *f*_*s*_(0) and *f*_*m*_(0) were set to one of four different values: period 45% shorter 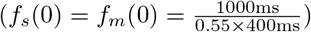, 30% shorter 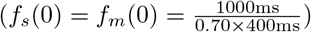, 15% shorter 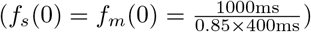, 15% longer 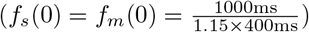, 30% longer 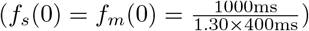, and 45% longer 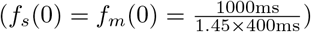 than ASHLE’s *f*_0_ period. In all simulations there was no external stimulus (i.e., *F* = 0). We analyzed the behavior of ASHLE for *γ* values of 0.01, 0.02, 0.04, 0.08. In each simulation, ASHLE oscillated for 50 seconds with initial conditions *z*_*s*_(0) = *z*_*m*_(0) = 0.001 + *i*0. Next, we found the location (in milliseconds) of the local maxima in the real-part of the oscillatory activity of *z*_*m*_. Then we found the difference between consecutive local maxima to obtain a sequence of IBIs. We calculated the linear regression between consecutive IBIs to obtain the resulting slope. Each line in Fig 8A shows the slopes obtained with different *γ* values and different initial conditions for *f*_*s*_(0) and *f*_*m*_(0). This analysis revealed that a value around *γ* = 0.02 will match the range of slope values observed in human data.

**Fig 8.**
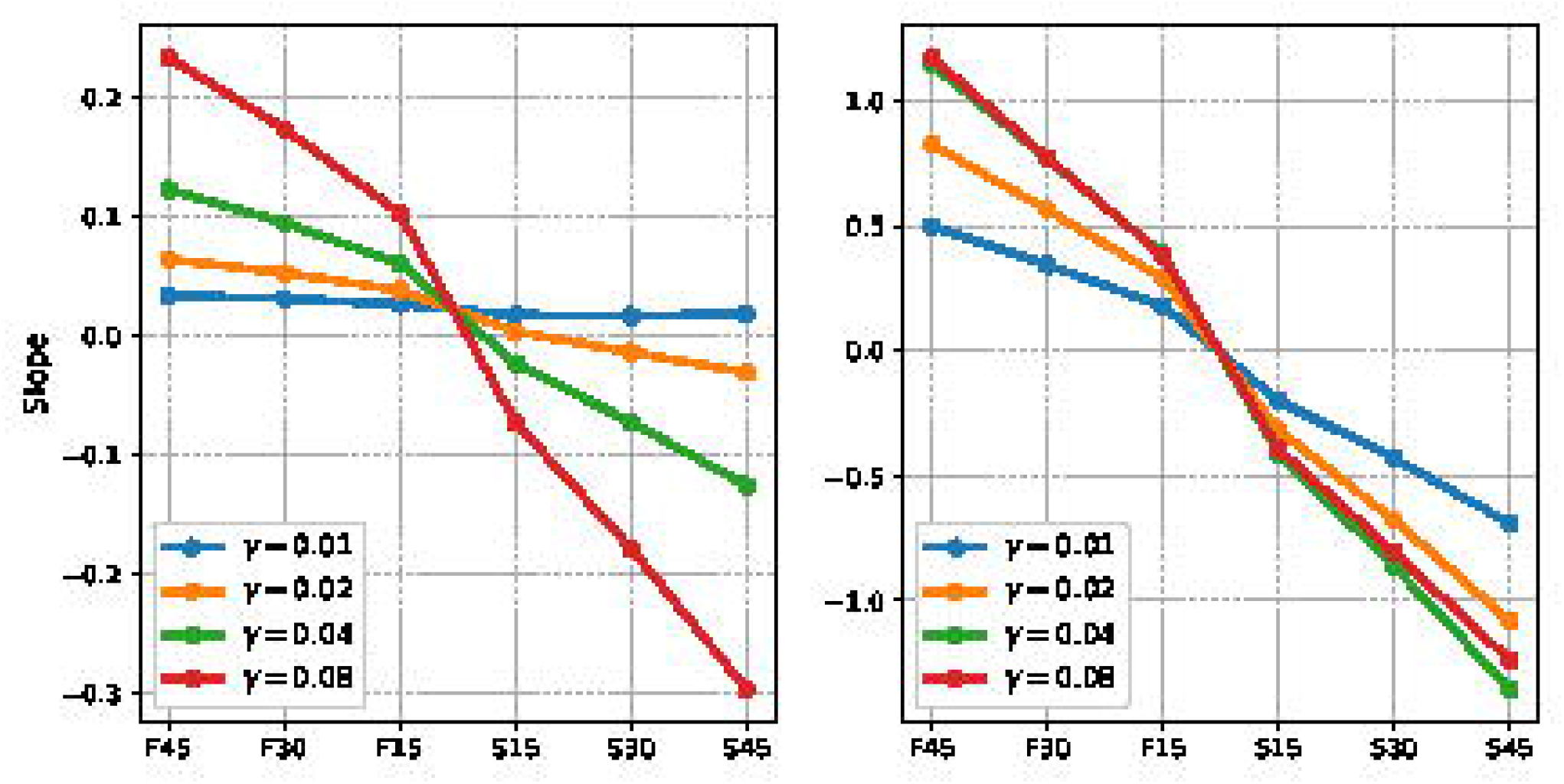
ASHLE slope values as a function of *γ* and initial frequency in the absence of a stimulus. (A) The effect of the *γ* parameter on the slope values between consecutive period lengths when ASHLE oscillates without a pacing stimulus, starting at a frequency that has a period 45% shorter 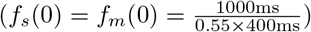, 30% shorter 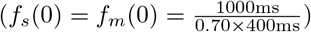, 15% shorter 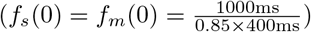, 15% longer 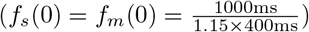, 30% longer 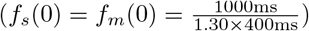, and 45% longer 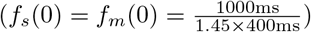 than ASHLE’s *f*_0_ period. (B) the same simulations but with an alternative single-oscillator model, showing a one-order-of-magnitude increase in the slope values

Finally, to determine if ASHLE really needs two oscillators, we also ran this analysis with an alternative single-oscillator model that consists of ASHLE’s Eq (3a) and (3b) where *f*_*m*_ is substituted by *f*_0_. Fig 8B shows the resulting slope values, revealing an order of magnitude increase in slope values. Such a single-oscillator model would need separate tuning and switching of the parameter pulling the oscillator to *f*_0_ to be able to explain the data in experiments 1 and 2. ASHLE, in contrast, natively shows two different time-scales depending on whether it is being stimulated or not, allowing it to explain both experiment 1 and 2 without the need for switching or tuning its parameters.

## Experiment 3: Duet musical performance between musicians with matching or mismatching SMTs

### Behavioral data for simulation

In another experiment, Zamm et al. [63] measured the SMP of 40 musicians and formed duets in two experimental groups: matching SMPs (*<*10ms IBI difference) and mismatching SMPs (*>*110ms IBI difference). There were 10 unique musician duets in each experimental group. Musician pairs were instructed to perform a simple unfamiliar melody together (16 beats in length), repeating the melody four consecutive times (64 beats total). When pairs of musicians performed the task, they first heard four metronome beats (400ms IOI) that established the common tempo. Experimenters measured the absolute asynchrony between each pair of synchronizing musicians throughout the entire performance. Because the same melody was repeated four times, they measured each melody repetition separately. Fig 4A shows their behavioral data. The mean absolute asynchrony between duets of musicians was larger when their SMPs did not match compared to when they matched.

### Setup, procedures and measurements

To obtain the musicians’ SMPs, we overlaid a grid over Figure 1 from their paper [63], which showed each musician’s measured SMP (in their original paper, the authors call the SMP as the “SPR”). This allowed us to precisely recover each musician’s SMP. We simulated 20 pairs of ASHLE models (10 pairs with similar natural frequencies and 10 pairs with dissimilar natural frequencies) synchronizing during 64 cycles. All simulations shared the same parameter values and initial conditions used in our experiment 1, except for the coupling strength between synchronizing ASHLE models (see model optimization in the next paragraph) and *f*_0_, which had a period that matched the SMP of a musician in the study. At the beginning of the simulation, two ASHLE models were stimulated by a complex-valued sinusoid *x*(*t*) = exp(*i*2*πf*_*x*_*t*) with a period of 400ms (*f*_*x*_ = 2.5). After these four cycles of sinusoidal stimulation, the stimulus stopped and the two ASHLE models stimulated each other with their respective *z*_*m*_. That is, in each duet simulation, after four cycles of sinusoidal stimulation, the input to the ASHLE No.1 was 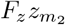 and the input to ASHLE No.2 was 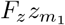, where *F*_*z*_ is the forcing strength between ASHLE models. After each simulation, we identified the location (in milliseconds) of local maxima in the real part of each ASHLE’s *z*_*m*_. Then, we measured the absolute asynchrony between the two synchronizing ASHLE model’s *z*_*m*_, obtaining 64 absolute asynchronies for each simulation. We divided these in four subsections (16 absolute asynchronies each) to simulate the four melody repetitions that pairs of musicians carried out, resulting in Fig 4B. We also simulated how different ASHLE models (with *f*_0_ periods linearly spaced between 350ms and 650ms) would carry out this task when synchronizing for 16 cycles with another ASHLE model with an *f*_0_ period difference of -220ms, -110ms, -10ms, 10ms, 110ms, and 220ms. Fig 4C shows these simulations, which are predictions of data that could be collected to test the accuracy of predictions made by ASHLE.

### Model optimization

There were two kinds of stimulation in this third experiment. During the first four cycles, similar to experiment 1, ASHLE was simulated by a sinusoid *x*(*t*) = *F* exp(*i*2*πf*_*x*_*t*) with a force *F* = 1 and *f*_*x*_ = 2.5. During the next 64 cycles, two ASHLE models synchronized with each other, so the input to the first ASHLE model is the second ASHLE model’s 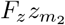 and the input to the second ASHLE model was the first ASHLE model’s 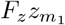. We found that using an *F*_*z*_ = 1 resulted in a lack of phase-locked synchronization between the two ASHLE, suggesting that *F*_*z*_ = 1 is too large and causes unstable dynamics between the two interacting ASHLE models. To improve stability, we reduced the value of *F*_*z*_ until we observed stable synchronization between all pairs of ASHLE models that we want to simulate, with the optimal *F*_*z*_ = 0.01. However, we also noted that the mean absolute asynchrony was considerably smaller between pairs of ASHLE models with similar *f*_0_ compared to the results for musician duets with matching SMTs. We believe that this difference was due to the lack of noise in our model. To improve our simulations we added gaussian noise to ASHLE’s *z*_*m*_ turning Eq (3c) into:

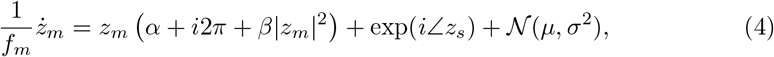

where *μ* = 0 is the mean and *σ* = 10 is the standard deviation of a normal distribution. Fig 4B shows the results obtained after we added the noise, which better approximate the behavioral data.

## Footnotes

1 Large et al. (2010) also included a *β*_2_ parameter that controls higher-order non-linear activity but we do not use it in the present study [28] (i.e. *β*_2_ = 0)

